# Alum:CpG adjuvant enables SARS-CoV-2 RBD-induced protection in aged mice and synergistic activation of human elder type 1 immunity

**DOI:** 10.1101/2021.05.20.444848

**Authors:** Etsuro Nanishi, Francesco Borriello, Timothy R. O’Meara, Marisa E. McGrath, Yoshine Saito, Robert E. Haupt, Hyuk-Soo Seo, Simon D. van Haren, Byron Brook, Jing Chen, Joann Diray-Arce, Simon Doss-Gollin, Maria De Leon, Katherine Chew, Manisha Menon, Kijun Song, Andrew Z. Xu, Timothy M. Caradonna, Jared Feldman, Blake M. Hauser, Aaron G. Schmidt, Amy C. Sherman, Lindsey R. Baden, Robert K. Ernst, Carly Dillen, Stuart M. Weston, Robert M. Johnson, Holly L. Hammond, Romana Mayer, Allen Burke, Maria E. Bottazzi, Peter J. Hotez, Ulrich Strych, Aiquan Chang, Jingyou Yu, Dan H. Barouch, Sirano Dhe-Paganon, Ivan Zanoni, Al Ozonoff, Matthew B. Frieman, Ofer Levy, David J. Dowling

**Affiliations:** Precision Vaccines Program, Division of Infectious Diseases, Boston Children’s Hospital, Boston, MA, USA; Department of Pediatrics, Harvard Medical School, Boston, MA, USA; Division of Immunology, Boston Children’s Hospital, Boston, MA, USA; Department of Microbiology and Immunology, University of Maryland School of Medicine, Baltimore, MD, USA; Department of Cancer Biology, Dana-Farber Cancer Institute, Boston, MA, USA; Department of Biological Chemistry and Molecular Pharmacology, Harvard Medical School, Boston, MA, USA; Research Computing Group, Boston Children’s Hospital, Boston, MA, USA; Ragon Institute of MGH, MIT, and Harvard, Cambridge, MA, USA.; Department of Microbiology, Harvard Medical School, Boston, MA, USA.; Department of Medicine, Brigham and Women’s Hospital, Boston, MA, USA; Department of Microbial Pathogenesis, University of Maryland School of Dentistry, Baltimore, MD, USA; Department of Pathology, University of Maryland Medical Center, Baltimore, MD, USA.; Texas Children’s Hospital Center for Vaccine Development, Baylor College of Medicine, Houston, TX, USA; National School of Tropical Medicine and Departments of Pediatrics and Molecular Virology & Microbiology, Baylor College of Medicine, Houston, TX, USA; National School of Tropical Medicine and Department of Pediatrics, Baylor College of Medicine, Houston, TX, USA; Center for Virology and Vaccine Research, Beth Israel Deaconess Medical Center, Harvard Medical School, Boston, MA, USA; Broad Institute of MIT & Harvard, Cambridge, MA, USA

## Abstract

Global deployment of vaccines that can provide protection across several age groups is still urgently needed to end the COVID-19 pandemic especially for low- and middle-income countries. While vaccines against SARS-CoV-2 based on mRNA and adenoviral-vector technologies have been rapidly developed, additional practical and scalable SARS-CoV-2 vaccines are needed to meet global demand. In this context, protein subunit vaccines formulated with appropriate adjuvants represent a promising approach to address this urgent need. Receptor-binding domain (RBD) is a key target of neutralizing antibodies (Abs) but is poorly immunogenic. We therefore compared pattern recognition receptor (PRR) agonists, including those activating STING, TLR3, TLR4 and TLR9, alone or formulated with aluminum hydroxide (AH), and benchmarked them to AS01B and AS03-like emulsion-based adjuvants for their potential to enhance RBD immunogenicity in young and aged mice. We found that the AH and CpG adjuvant formulation (AH:CpG) demonstrated the highest enhancement of anti-RBD neutralizing Ab titers in both age groups (∼80-fold over AH), and protected aged mice from the SARS-CoV-2 challenge. Notably, AH:CpG-adjuvanted RBD vaccine elicited neutralizing Abs against both wild-type SARS-CoV-2 and B.1.351 variant at serum concentrations comparable to those induced by the authorized mRNA BNT162b2 vaccine. AH:CpG induced similar cytokine and chemokine gene enrichment patterns in the draining lymph nodes of both young adult and aged mice and synergistically enhanced cytokine and chemokine production in human young adult and elderly mononuclear cells. These data support further development of AH:CpG-adjuvanted RBD as an affordable vaccine that may be effective across multiple age groups.

**One Sentence Summary:** Alum and CpG enhance SARS-CoV-2 RBD protective immunity, variant neutralization in aged mice and Th1-polarizing cytokine production by human elder leukocytes.

## INTRODUCTION

The coronavirus disease 2019 (COVID-19) pandemic caused by severe acute respiratory syndrome coronavirus 2 (SARS-CoV-2) resulted in a serious threat to humanity. Rapid deployment of safe and effective vaccines is proving key to reducing morbidity and mortality of COVID-19, especially in high-risk populations such as the older adults (*1*). Novel vaccine technologies including mRNA and adenoviral vector vaccines have dramatically accelerated the process of vaccine development, shown high efficacy in preclinical and clinical studies, and therefore been granted Emergency Use Authorization by the Food and Drug Administration (*2-9*). Unfortunately, worldwide access to these vaccines may be limited by the need for ultra-cold storage (mRNA vaccines), cost, and concerns regarding global scalability especially in the third world (*1*). This situation not only represents a major ethical problem but may also promote the emergence of vaccine-resistant SARS-CoV-2 strains due to high infection rates in unvaccinated regions (*10*). Thus, ongoing efforts are needed to investigate additional affordable, easily scalable, and effective vaccine approaches against SARS-CoV-2 to improve global access. To this end, alternative platforms such as inactivated and protein subunit SARS-CoV-2 vaccines have entered different stages of clinical development and in some cases have already been deployed at the population level (*11-17*). These approaches may play an essential role in the global fight against COVID-19 since they utilize well-established technologies, do not require low temperature storage, and have proven safety and effectiveness in various age groups including young children and the elderly.

With the exception of inactivated viruses, most SARS-CoV-2 vaccine candidates aim to target the SARS-CoV-2 Spike glycoprotein, as it is required for binding to the human receptor angiotensin-converting enzyme 2 (ACE2) and subsequent cell fusion. In particular, the receptor-binding domain (RBD) of the Spike protein plays a key role in ACE2 binding and is targeted by many neutralizing antibodies (Abs) that exert a protective role against SARS-CoV-2 infection (*18-20*). RBD is an attractive candidate for a SARS-CoV-2 subunit vaccine and is relatively easy to produce at scale (*21, 22*); however, it is poorly immunogenic on its own. Structural biology-based vaccine design has been employed to overcome this limitation and has generated encouraging results in preclinical and clinical studies (*22-29*). A complementary approach to increase the immunogenicity of vaccine antigens consists of using adjuvants, which can enhance antigen immunogenicity by activating receptors of the innate immune system called pattern-recognition receptors (PRRs) and/or modulating antigen pharmacokinetics (*30, 31*). Adjuvant formulations of aluminum salts and PRR agonists enhance vaccine immune responses compared to aluminum salts or PRR agonists alone (*32*). AS04 was the first adjuvant system composed of aluminum salts and a PRR agonist, specifically the TLR4 agonist monophosphoryl lipid A (MPLA), to be included in a licensed human papillomavirus and hepatitis B vaccines (*32*). Thus, combinations of aluminum salts and PRR agonists represent a promising adjuvant platform to enhance RBD immunogenicity.

Here, we evaluated several combinations of PRR agonists and aluminum hydroxide (AH) and found that the TLR9 agonists CpG oligodeoxynucleotides formulated with AH and RBD dramatically enhanced immune response towards RBD in young mice using a prime-boost immunization schedule. The AH:CpG-adjuvanted RBD vaccine also elicited a robust anti-RBD immune response in aged mice, with the administration of an additional boost dose generating an anti-RBD Ab response comparable to young adult mice and providing complete protection from live SARS-CoV-2 challenge. Overall, our comprehensive, head-to-head adjuvant comparison study demonstrates that AH:CpG co-adjuvantation can overcome both the poor immunogenicity of RBD and immunosenescence, supporting this approach for development of a scalable, affordable, and safe global SARS-CoV-2 vaccine tailored for older adults.

## RESULTS

### Evaluation of multiple AH:PRR agonist formulations in young adult mice

We first evaluated whether distinct AH:PRR agonist formulations can overcome the low immunogenicity of monomeric RBD proteins. To this end, we performed a comprehensive comparison of PRR agonists, including 2’3’-cGAMP (stimulator of IFN genes (STING) ligand), Poly (I:C) (TLR3 ligand), PHAD (synthetic MPLA, TLR4 ligand), and CpG-ODN 2395 (TLR9 ligand). Each PRR agonist was formulated with and without AH. We also included AS01B (a liposome-based adjuvant containing MPLA and the saponin QS-21) as a clinical-grade benchmark adjuvant with potent immunostimulatory activity. The immunogenicity of vaccine formulations was first evaluated in 3-month-old young adult mice. Mice were immunized intramuscularly twice with 10 µg of monomeric RBD protein formulated with or without adjuvant, in a two-dose prime-boost regimen (Days 0 and 14). Two weeks after the boost immunization, humoral immune responses were evaluated. AH:PRR agonist formulations enhanced both anti-RBD Ab titers and inhibition of RBD binding to human ACE2 (hACE2) as compared to their respective non-AH adjuvanted formulations (**Fig 1A-C**). The Ab response elicited by AH alone was highly skewed to IgG1, with minimal inhibition of hACE2/RBD binding (**Fig 1D, E**). Among various AH:PRR agonist formulations, AH:CpG demonstrated the highest induction of total IgG, IgG1, and IgG2a along with a balanced IgG2a/IgG1 ratio (**Fig 1A-D**). Furthermore, the AH:CpG formulation significantly enhanced hACE2/RBD binding inhibition compared to all the other AH:PRR agonist formulations (**Fig 1E**). Abs induced by monomeric RBD immunization recognized the native trimeric Spike protein, as demonstrated by a binding ELISA with prefusion stabilized form of spike trimer (**Fig 1F**). To assess long-term immunogenicity, we then evaluated Ab responses and hACE2/RBD binding inhibition on Day 210 (**Fig 1G-J**). Of note, AH:CpG formulation maintained high hACE2/RBD binding inhibition while other adjuvant formulations waned their immune responses (**Fig 1E, J**).

**Figure 1.**
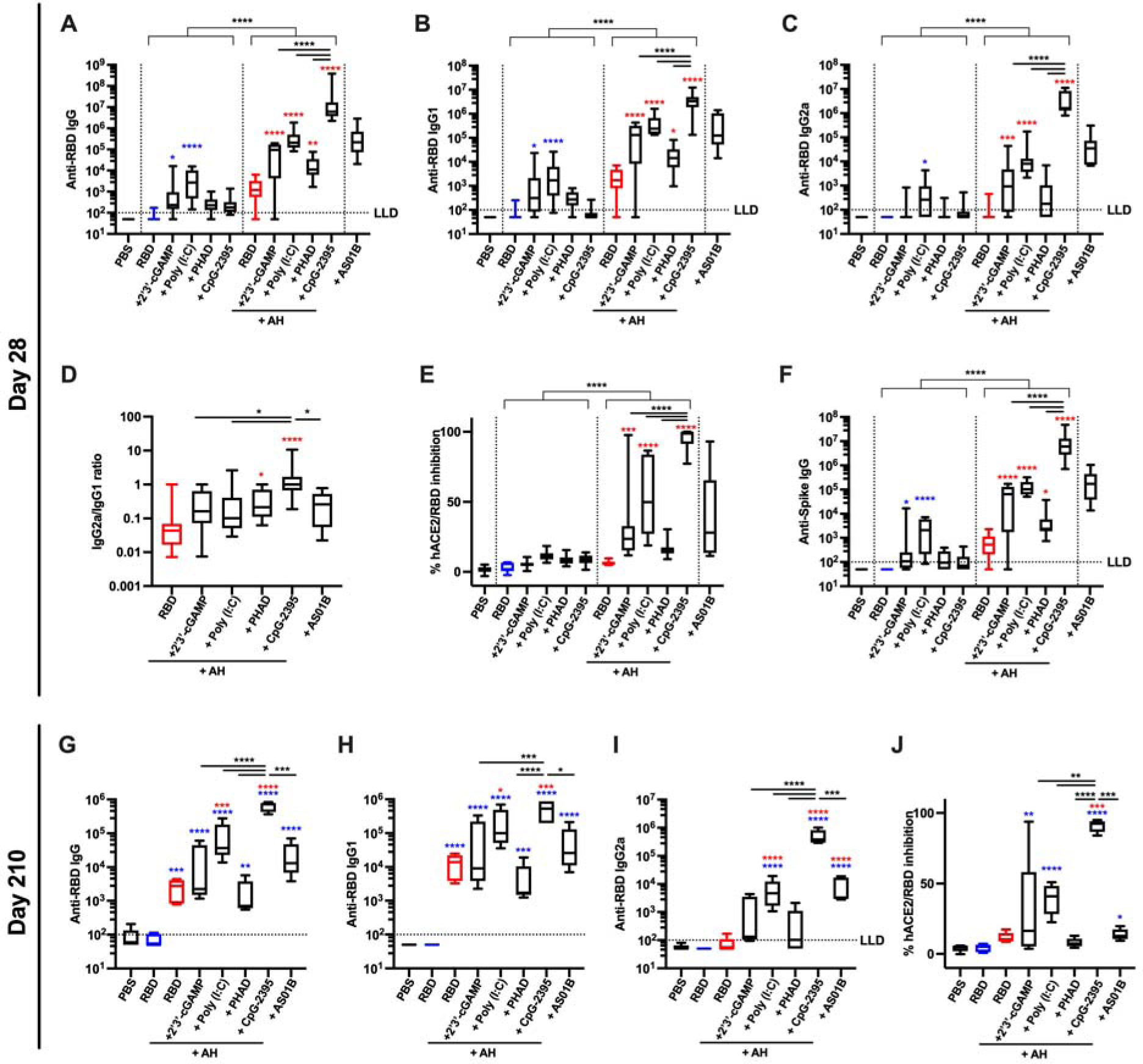
RBD formulated with AH:CpG induces robust production of anti-RBD neutralizing antibodies in young adult mice. Young adult, 3-month-old BALB/c mice were immunized IM on Days 0 and 14 with 10 µg of monomeric SARS-CoV-2 RBD protein with indicated adjuvants. Each PRR agonist was administered alone or formulated with aluminum hydroxide (AH). (**A–F**) Serum samples were collected on Day 28, and (**A**) Anti-RBD IgG, (**B**) IgG1, (**C**) IgG2a, (**D**) IgG2a/IgG1 ratio, (**E**) hACE2/RBD inhibition rate, and (**F**) anti-Spike IgG were assessed. N=10 per group. Data were combined from two individual experiments. (**G–J**) Serum samples were collected on Day 210, and (**G**) Anti-RBD IgG, (**H**) IgG1, (**I**) IgG2a and (**J**) hACE2/RBD inhibition rate were assessed. N=5 per group. Data were analyzed by two-way (**A–C, E–F**) (AH and PRR agonist) or one-way (**D, G–J**) ANOVAs followed by post-hoc Tukey’s test for multiple comparisons. \**P* <0.05, \*\**P* <0.01, \*\*\**P* <0.001, **** *P* <0.0001. Blue and red colored asterisks respectively indicate comparisons to RBD and AH adjuvanted RBD groups. Box-and-whisker plots represent the minimum, first quartile, median, third quartile, and maximum value. LLD, lower limit of detection.

### AH:CpG-formulated RBD vaccine is immunogenic in aged mice

To assess the vaccine response in the context of aging, the immunogenicity of RBD vaccines adjuvanted with AH:PRR agonists was further studied in aged mice (14-month-old). Similar to young mice, the AH:CpG formulation also elicited the highest humoral immune response after prime-boost immunization in aged mice (**Fig 2A-F**). Of note, the vaccine adjuvanted with AH:CpG produced significantly higher hACE2/RBD inhibition and neutralizing titers compared to the vaccine adjuvanted with AS01B, which is known as a potent adjuvant in the human elderly population (*33, 34*) (**Fig 2E, F**). However, Ab levels were generally lower in aged mice, and the magnitude of the immune response of aged mice receiving the AH:CpG vaccine was significantly lower than that of young mice, suggesting an impaired vaccine response due to immunosenescence in the elderly population (**Fig S1**). To determine whether an additional dose can improve vaccine immunogenicity in aged mice, we administered a second booster dose two weeks after the last immunization. On Day 42 (two weeks after the 2^nd^ boost), enhancement in humoral responses was observed in AH:PRR agonist formulations (**Fig 2G-L**). Notably, a significant enhancement of hACE2/RBD inhibition was observed in aged mice receiving the two-boost AH:CpG vaccination regimen, with inhibition reaching the level of young mice that had received AH:CpG in a prime-boost regimen (**Fig S1**). High serum neutralizing Ab titers were observed in the AH:CpG and AS01B adjuvanted groups after the 2^nd^ boost but not in the non-adjuvanted nor AH alone-adjuvanted RBD groups. Assessment of cytokine production by splenocytes isolated from immunized mice and restimulated *in vitro* with Spike peptides demonstrated high Th1 (IFNγ and IL-2) and low Th2 (IL-4) cytokine production in the AH:CpG and AS01B groups (**Fig 2M**). These results demonstrate that the AH:CpG-adjuvanted RBD vaccine is highly immunogenic in aged mice and an additional booster dose can further enhance anti-RBD humoral responses to match those observed in young mice.

**Figure 2.**
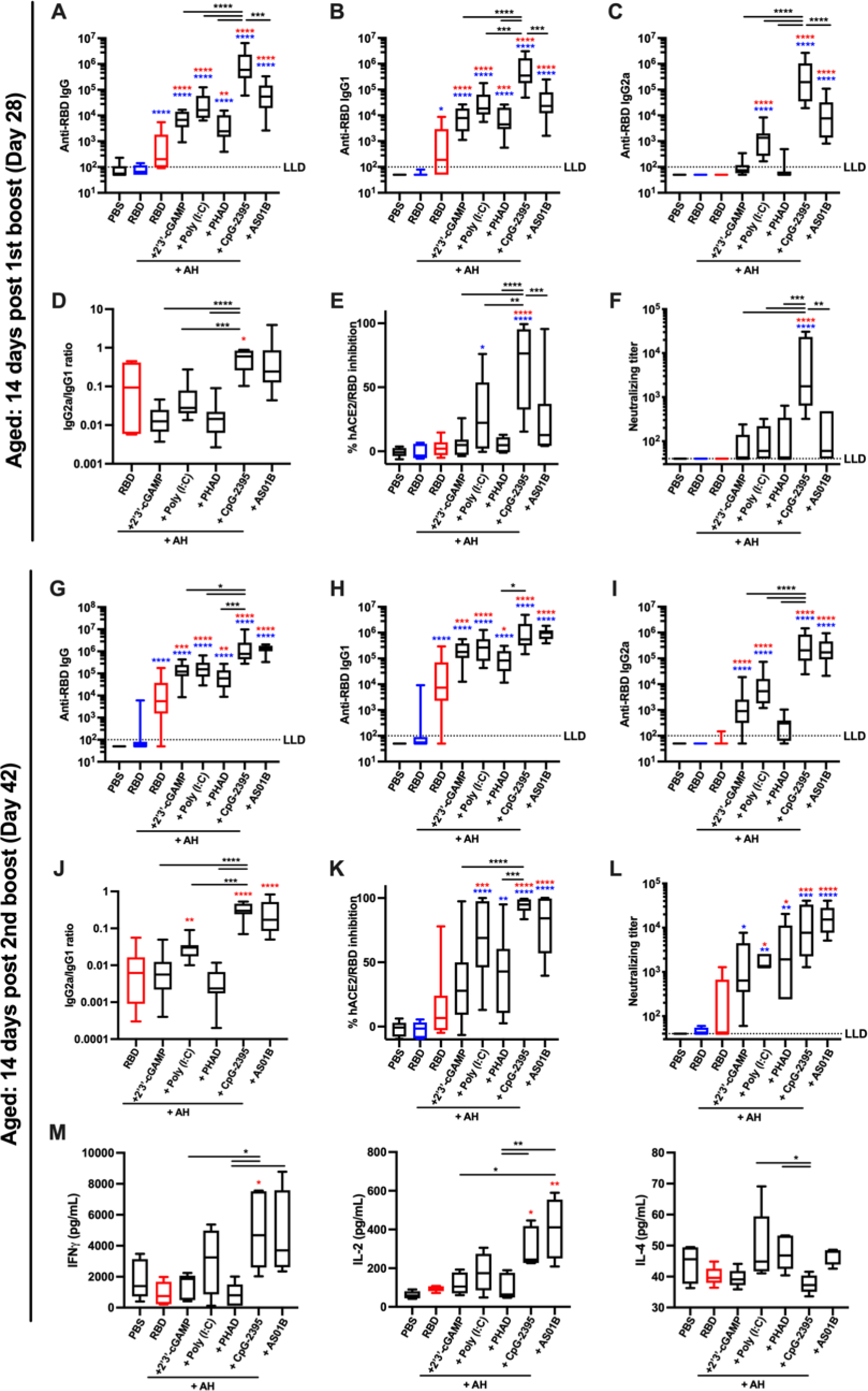
AH:CpG adjuvant formulation elicits a robust anti-RBD response in aged mice. Aged, 14-month-old BALB/c mice were immunized IM on Days 0, 14, and 28 with 10 µg of monomeric SARS-CoV-2 RBD protein with indicated adjuvants. Each PRR agonist was formulated with aluminum hydroxide (AH). Serum samples were collected and analyzed on day 28 prior to the 2nd boost (**A–F**), and day 42 (**G–L**). (**A, G**) Anti-RBD IgG, (**B, H**) IgG1, (**C, I**) IgG2a, (**D, J**) IgG2a/IgG1 ratio, (**E, K**) hACE2/RBD inhibition rate, and (**F, L**) neutralizing titer were assessed. N=9–10 per group. Data were combined from two individual experiments and analyzed by one-way ANOVAs followed by post-hoc Tukey’s test for multiple comparisons. (**M**) Splenocytes were collected 2 weeks after the final immunization and stimulated with a SARS-CoV 2 Spike peptide pool in the presence of anti-CD28 antibody (1 μg/mL). After 24 (for IL-2 and IL-4) and 96 (for IFNγ) hours, supernatants were harvested and cytokine levels were measured by ELISA. N=4-5 per group. Data were log-transformed and analyzed by one-way ANOVAs followed by post-hoc Tukey’s test for multiple comparisons. \**P* <0.05, \*\**P* <0.01, \*\*\**P* <0.001, **** *P* <0.0001. Blue and red colored asterisks respectively indicate comparisons to RBD and AH adjuvanted RBD groups. Box-and-whisker plots represent the minimum, first quartile, median, third quartile, and maximum value. LLD, lower limit of detection.

### AH:CpG-formulated RBD vaccine protects aged mice from lethal viral challenge

Neutralizing Abs are key to protecting from SARS-CoV-2 infection. Since RBD formulated with AH:CpG elicited high titers of neutralizing Abs, we assessed the protection of immunized mice in a challenge model. To this end, we employed the mouse-adapted SARS-CoV-2 MA10 virus strain (*35*). When tested in young (3-month-old) and aged (14-month-old) BALB/c mice, SARS-CoV-2 MA10 elicited dose-dependent weight loss (**Fig 3A, B**). Notably, aged mice challenged with 10^3^ PFU or more exhibited dose-dependent mortality by 4 days post-infection (dpi) (**Fig 3C**). None of the young mice died by 4 dpi, including those that received the highest viral dose, in contrast with aged mice. Next, immunized aged mice were challenged with SARS-CoV-2 MA10 six weeks after the second boost. Bodyweight changes were assessed daily up to 4 dpi when the mice were sacrificed for viral titer and histopathology analyses. Aged mice immunized with the AH:CpG and AS01B adjuvanted vaccines showed no weight loss up to 4 dpi, whereas aged mice immunized with non-adjuvanted or AH-adjuvanted RBD showed rapid and significant bodyweight loss of >10% through 4 dpi (**Fig 4A**). Lung tissues were harvested and tested for SARS-CoV-2 viral titer in lung. No detectable live virus in lung tissues was observed in the AH:CpG and AS01B adjuvanted groups, while viral titers were detectable in the vehicle, non-adjuvanted, and AH-adjuvanted groups (**Fig 4B**). Histopathological analysis conducted in lung tissues further confirmed the reduced SARS-CoV-2 infection in aged animals vaccinated with AH:CpG and AS01B adjuvants (**Fig 4C, D**).

**Figure 3.**
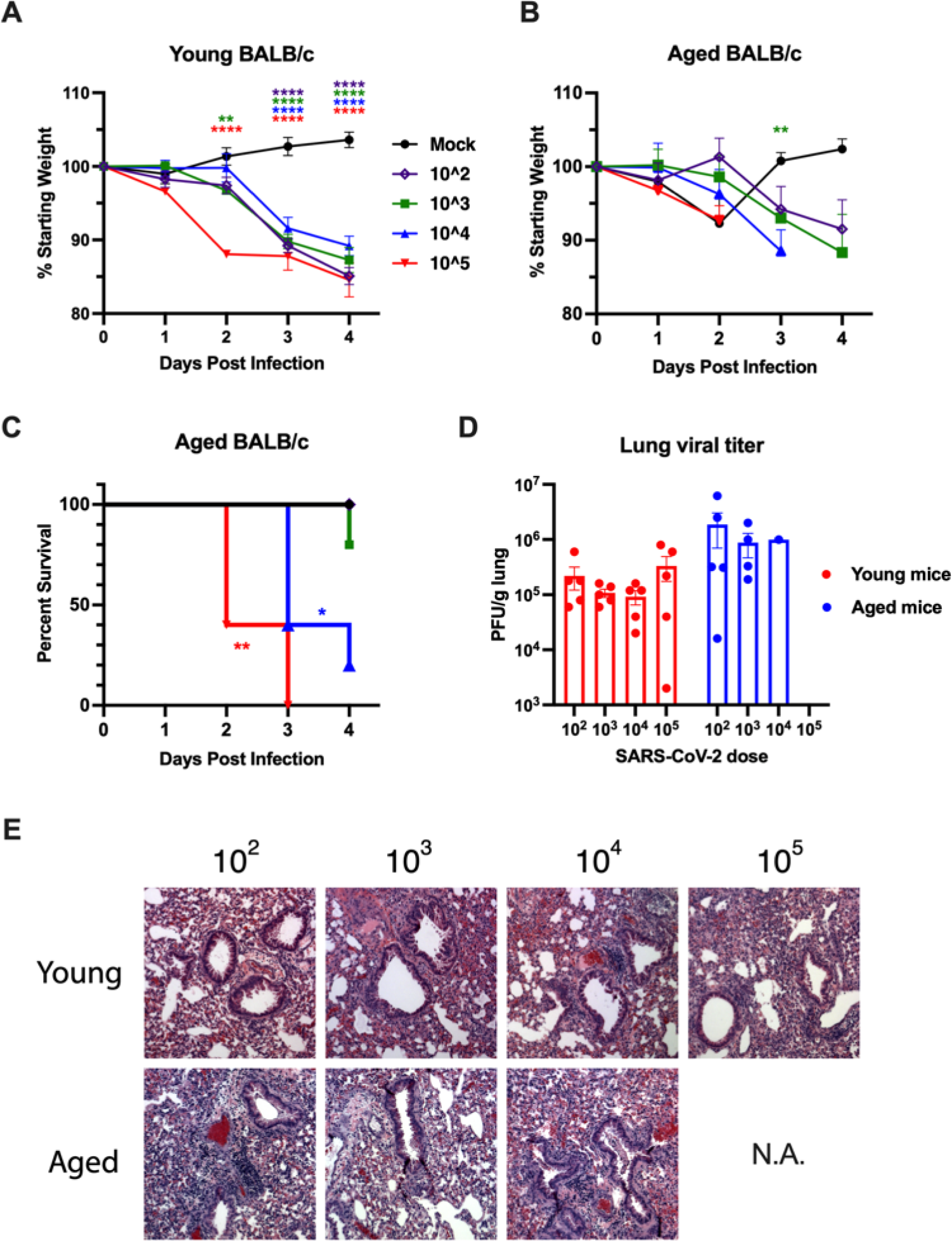
SARS-CoV-2 challenge model of young and aged mouse recapitulates human age-specific pathogenesis. Young (3-month-old) and aged (14-month-old) naïve BALB/c mice were challenged IN with mock (PBS), or 10^2^, 10^3^, 10^4^, and 10^5^ PFU of mouse-adapted SARS-CoV-2 (MA10). Bodyweight change of (**A**) young adult and (**B**) aged mice were assessed daily up to 4 days post infection. Data represent mean and SEM with body weights only shown for surviving mice at each time-point. Data were analyzed by one-way ANOVA followed by Dunnett’s test for comparisons against the PBS group. (**C**) Survival rate of aged mice. Data were analyzed by log-rank test in comparison to PBS group. (**D**) Viral titer in lung homogenates at 4-days post SARS-CoV-2 challenge (young: n=5 per group, aged: n=5 for 10^2^; n=4 for 10^3^; n=1 for 10^4^; and n=0 for 10^5^). Results represent mean ± SEM. (**E**) Representative lung histological images at 4-days post challenge. H&E is shown. \**P* <0.05, \*\**P* <0.01, \*\*\**P* <0.001, **** *P* <0.0001.

**Figure 4.**
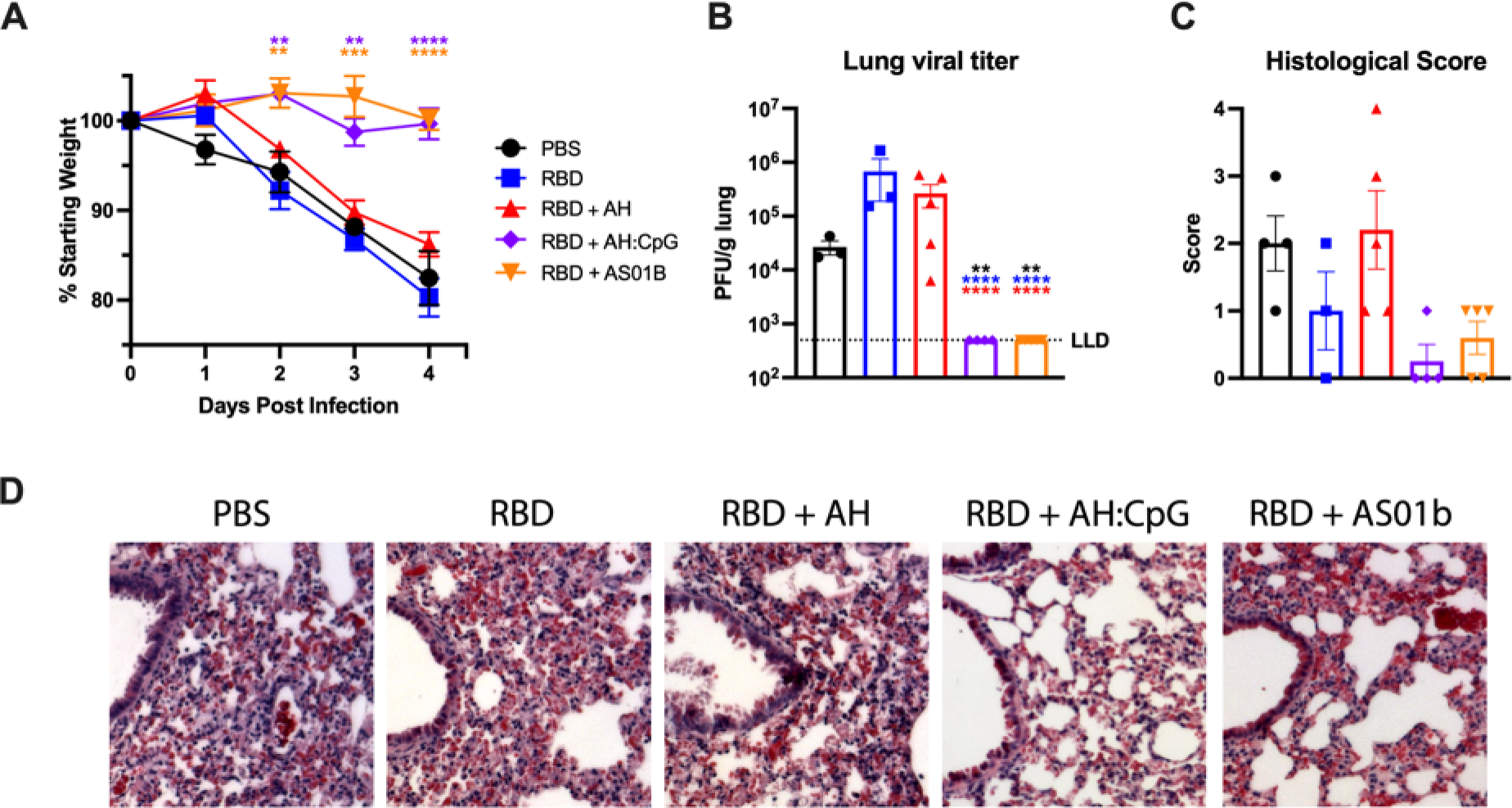
AH:CpG-adjuvanted vaccine protects aged mice from SARS-CoV-2 challenge. Aged, 14-month-old BALB/c mice were immunized as in Figure 2. On Day 70 (6 weeks post 2^nd^ boost), mice were challenged IN with 10^3^ PFU of mouse-adapted SARS-CoV-2 (MA10). (**A**) Bodyweight changes were assessed daily up to 4 days post infection. Data represent mean and SEM with body weights shown for surviving mice at each time-point (one mouse in RBD group died at 4 days post infection). Data were analyzed by one-way ANOVA followed by Dunnett’s Test for comparisons between PBS group. (**B**) Viral titer in lung homogenates at 4-days post SARS-CoV-2 challenge. Results represent mean ± SEM. Data were analyzed by one-way ANOVA followed by post-hoc Tukey’s test for multiple comparisons. \*\**P* <0.01, **** *P* <0.0001. Black, blue and red colored asterisks respectively indicate comparisons to PBS, RBD, and RBD + aluminum hydroxide (AH) groups. LLD, lower limit of detection. (**C**) Lung interstitial inflammation was evaluated and converted to a score of 0-4 with 0 being no inflammation and 4 being most severe. (**D**) Representative lung histological images at 4-days post challenge. H&E is shown. N=4-5 animals per group.

### AH:CpG-formulated RBD and Spike mRNA vaccines elicit comparable levels of neutralizing antibodies against wild type SARS-CoV-2 and variants

Recently, it has been reported that SARS-CoV-2 mRNA vaccines are more immunogenic than RBD adjuvanted with oil-in-water emulsions (*36*). To assess whether this is a general feature of RBD protein vaccines, we used the clinical-grade authorized BNT162b2 Spike mRNA vaccine (Pfizer-BioNTech) as a benchmark and compared it to RBD formulated with AddaS03 (a commercially available version of the oil-in-water emulsion AS03) and to AH:CpG in aged mice. Along with CpG-2395, we also tested CpG-1018, which is included in the Heplisav-B vaccine and has also been tested in combination with Spike/RBD and AH in SARS-CoV-2 studies including human vaccine trials (*12, 16, 37*). In accordance with previously published data, the mRNA vaccine was highly immunogenic, while RBD formulated with AddaS03 failed to induce significant levels of neutralizing Abs (**Fig 5A-D**). Of note, both AH:CpG formulations elicited levels of anti-RBD (**Fig 5A**), anti-Spike (**Fig 5B**) and neutralizing Abs (**Fig 5C, D**) comparable to or greater than the mRNA vaccine.

**Figure 5.**
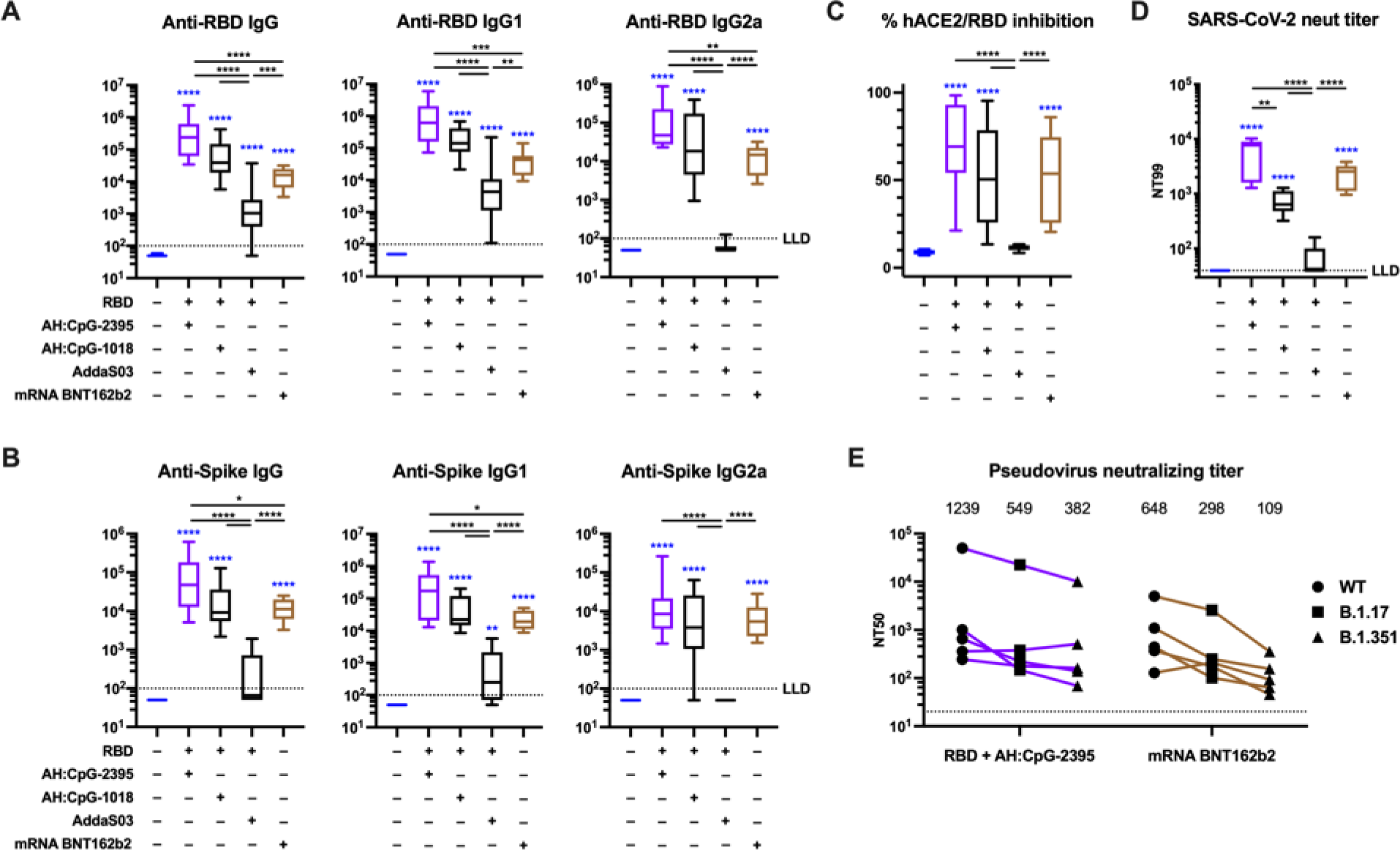
AH:CpG-adjuvanted RBD vaccines and an authorized spike mRNA vaccine elicit comparable levels of neutralizing antibodies in aged mice. Aged, 14-month-old BALB/c mice were immunized IM on Days 0 and 14 with monomeric SARS-CoV-2 RBD protein with indicated adjuvants, or BNT162b2 Spike mRNA vaccine as described in Methods. Serum samples were collected and analyzed on Day 28. (**A**) Anti-RBD binding ELISA, (**B**) anti-Spike binding ELISA, (**C**) hACE2/RBD inhibition rate, and (**D**) SARS-CoV-2 virus neutralizing titer were assessed. N=9–10 per group. Data were combined from two individual experiments and analyzed by one-way ANOVAs followed by post-hoc Tukey’s test for multiple comparisons. (**E**) Pseudovirus neutralizing titers against wild-type or the B.1.17 or B.1.351 variants were assessed. N=5 per group. The numbers indicate GMT. Each symbol represents an animal. \**P* <0.05, \*\**P* <0.01, \*\*\**P* <0.001, **** *P* <0.0001. Blue colored asterisks indicate comparisons to PBS group. Box-and-whisker plots represent the minimum, first quartile, median, third quartile, and maximum value. LLD, lower limit of detection.

SARS-CoV-2 variants such as B.1.1.7 and B.1.351 have emerged with reduced neutralization from serum samples of convalescent or vaccinated individuals (*38-41*). A recent report showed that the mRNA BNT162b2 vaccine maintained its effectiveness against severe COVID-19 with the B.1.351 variant at greater than 90% (*42*). We therefore evaluated whether RBD + AH:CpG and mRNA BNT162b2 vaccines elicit neutralizing Abs against these variants. As expected, we observed reduced titers against the variants, especially against the B.1.351 (**Fig 5E**). The neutralization titers of RBD + AH:CpG decreased by 3.2-fold against B.1.351, and the mRNA BNT 162b2 decreased by 6.0-fold. Neutralizing titers against the B.1.351 were comparable between RBD + AH:CpG (GMT 382) and mRNA BNT162b2 (GMT 109).

### Innate signaling potentiated by AH:CpG formulation is well preserved in aged mice

Lymph nodes (LNs) are critical sites for the interaction between innate and adaptive immune systems and orchestrate the development of vaccine immune responses (*43, 44*). Specifically, activation of the innate immune system can induce a rapid response in the LN characterized by LN expansion, which is driven by lymphocyte accrual and expression of proinflammatory molecules (*45, 46*). To gain further insights into the mechanism of action of the AH:CpG formulation, we collected draining LNs (dLNs) 24 hours post injection of AH:CpG or either adjuvant alone. CpG and AH:CpG induced comparable dLN expansion in both age groups (**Fig 6A**). To characterize the molecular events associated with these treatments further, RNA isolated from dLNs after injection of vehicle, CpG, or AH:CpG was subjected to a quantitative real-time PCR array comprised of 157 genes related to cytokines, chemokines, and type 1 IFN responses. Principal component analysis and hierarchical cluster analysis demonstrated a marked separation between AH and CpG-containing treatments, whereas similar patterns were observed between groups treated with AH:CpG and CpG alone in both age groups (**Fig 6B, C**). Generalized linear model analysis comparing gene expressions after AH, CpG, and AH:CpG treatments further revealed similar gene enrichment patterns between young adult and aged mice (**Fig 6D, E**). These results suggest that CpG and AH:CpG activate similar pathways in young and aged mice to elicit a LN innate response.

**Figure 6.**
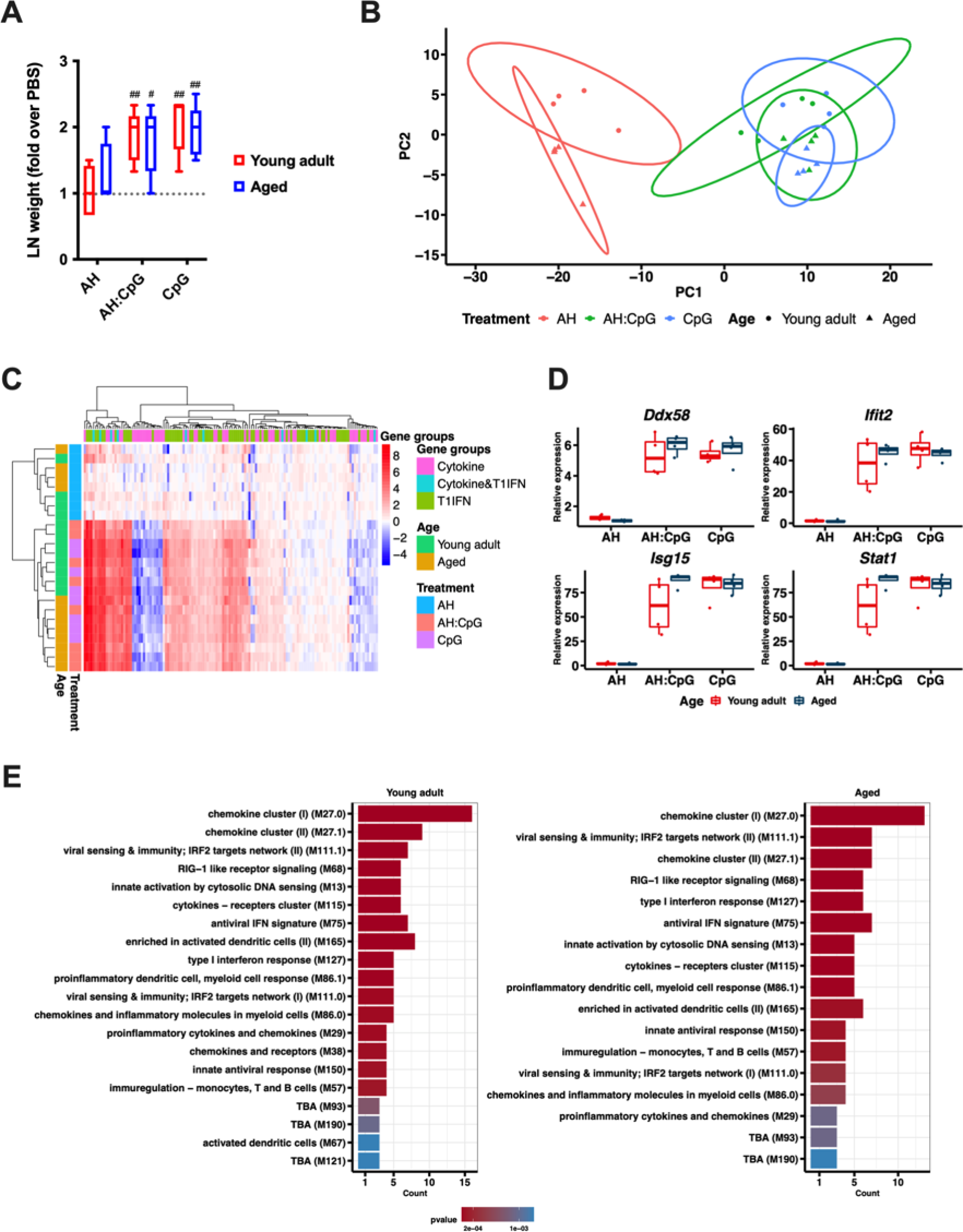
AH:CpG elicits comparable lymph node innate responses in young and aged mice. Young (3-month-old) and aged (14-month-old) mice were subcutaneously injected with aluminum hydroxide (AH), CpG, or AH:CpG. 24 hours later, draining lymph nodes (dLNs) were collected and RNA was extracted. (**A**) Weights of dLNs were measured and expressed as fold over contralateral, PBS-injected LN. N=5 per group. # and ## respectively indicate *P* <0.05 and 0.01 when comparing each group against the value 1 (which represents the contralateral control sample expressed as fold). (**B–E**) RNA isolated from dLNs was subjected to a quantitative real-time PCR array comprised of 157 genes related to cytokines, chemokines, and type 1 IFN responses. N=4 animals per group. (**B**) Principal component analysis demonstrated a marked separation by treatment and age. (**C**) Unsupervised hierarchical clustering revealed major differences between treatments and highlighted the marked difference between AH and CpG-containing treatments. Each column represents gene categories and rows represent samples. (**D**) Generalized linear model comparing treatment and age with each gene was performed. The top 4 significant genes (*Ddx58*, *Ifit2*, *Isg15*, *Stat1*) were selected and plotted with their relative expression values by age and treatment. Statistical analysis of the plots employed the Kruskal-Wallis test to compare mean differences across groups and Wilcoxon test to compare between ages. (**E**) Enrichment analysis of differentially expressed genes using the blood transcriptional modules (Li et al., 2013-PMC: 24336226) was performed from the significant gene results after the generalized linear model by treatment. The top 20 modules are summarized per age.

### AH:CpG synergistically enhances proinflammatory cytokines from human elderly PBMCs

In order to assess the translational relevance of an adjuvant formulation it is key to confirm its ability to activate human immune cells. To this end, we stimulated human peripheral blood mononuclear cells (PBMCs) isolated from young adults (18-40 years old) and elder adults (≥ 65 years old) with CpG, AH, and the admixed AH:CpG formulation and measured cytokine and chemokine production. Whereas AH induced limited or no cytokine production, both CpG alone and AH:CpG activated young adult and elderly PBMCs in a concentration-dependent manner (**Fig 7A-D**, **Fig S2**). PBMCs of both age groups treated with AH:CpG produced significantly higher levels of various proinflammatory cytokines and chemokines than those treated with CpG alone (**Fig 7A-D**). Of note, CpG and AH synergistically induced IL-6, IL-10, TNF, CCL3, and GM-CSF production in both young adult and elderly PBMCs, as defined mathematically (D value, see Methods) (**Fig 7C, D**, **Fig S2**).

**Figure 7.**
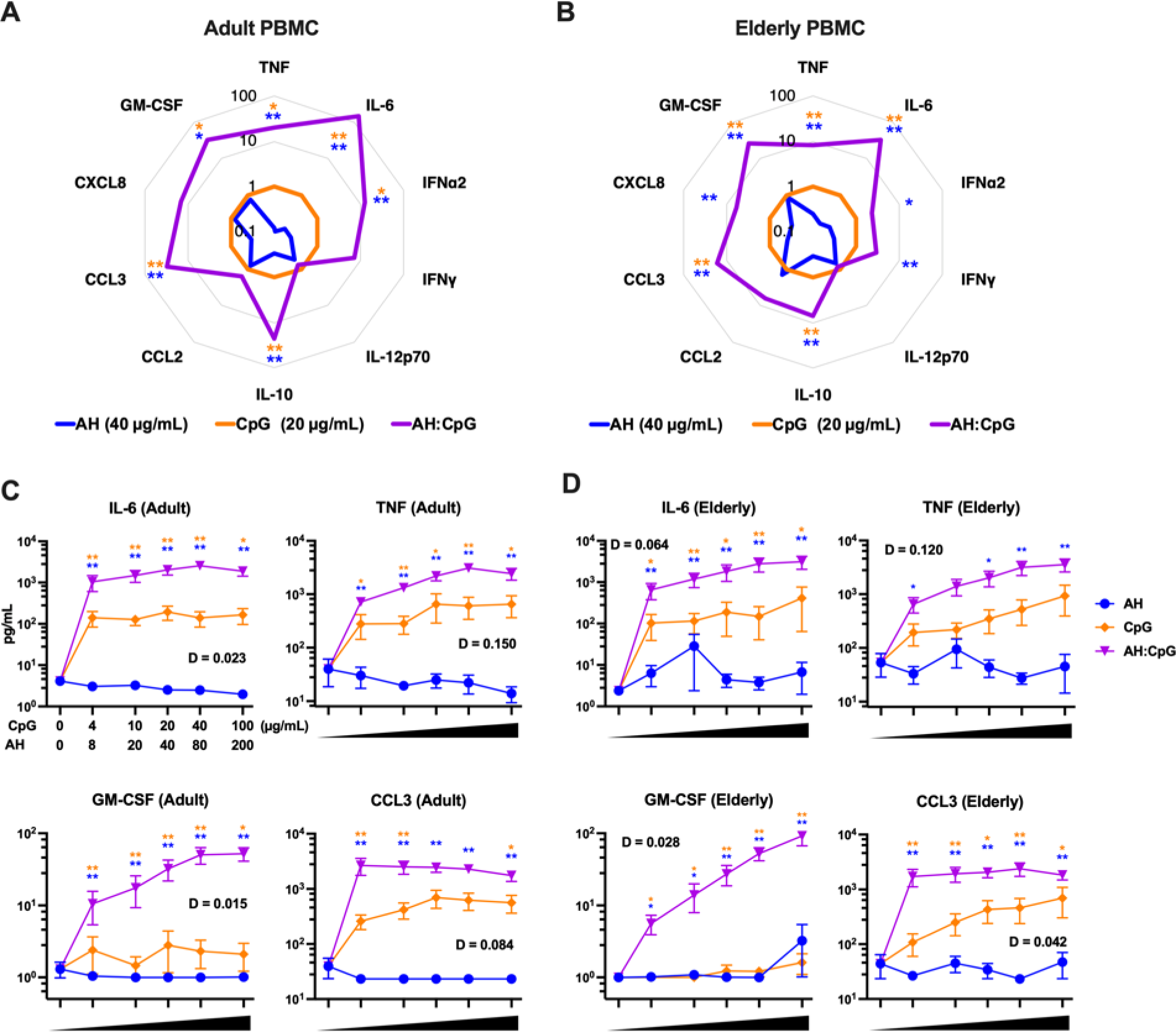
AH:CpG synergistically enhances proinflammatory cytokine production from human adult and elderly PBMCs. Human PBMCs collected from young adult (**A, C**) and elderly individuals (**B, D**) were cultured *in vitro* for 24 h with CpG alone (4, 10, 20, 40, and 100 μg/mL), aluminum hydroxide (AH) alone (8, 20, 40, 80, and 200 μg/mL), or a combination of both. Supernatants were collected for multiplexing bead array. N=6 per age group. (**A-B**) Radar plot analysis of cytokines and chemokines are presented as a fold-change over the CpG alone group for the 20 µg/mL CpG and 40 µg/mL AH conditions. (**C-D**) Results represent mean ± SEM. Unpaired Mann-Whitney tests were applied at each concentration. Blue and yellow colored asterisks indicate comparisons of AH:CpG formulation to AH and CpG alone groups, respectively. \**P* <0.05, \*\**P* <0.01. Level of synergy was calculated using an adapted Loewe definition of additivity (D <1: synergy, D=1: additivity, D >1: antagonism).

## DISCUSSION

The risk of COVID-19-related morbidity and mortality increases with age (*47, 48*). Currently authorized SARS-CoV-2 vaccines have proven effective at preventing severe COVID-19 (*2-4*). Nevertheless, there is still the need to develop affordable and accessible vaccines that can provide protection across several age groups, especially for low- and middle-income countries (*1, 10, 49*). Protein subunit vaccines formulated with appropriate adjuvants represent a promising strategy to address this urgent need. Here, we performed a comprehensive head-to-head comparison of multiple adjuvants in age-specific *in vivo* and *ex vivo* animal models, along with age-specific human *in vitro* screening, to determine the appropriate adjuvant for a SARS-CoV-2 RBD vaccine in the young and the aged, focusing on the innate and humoral immune response reported to align best with known correlates of protection (*50, 51*). We found that the AH:CpG adjuvant formulation enhances anti-RBD neutralizing Ab titers and type 1 immunity (i.e. IgG2a switching, Th1 polarization) in both age groups. Aged mice immunized with AH:CpG are protected from live SARS-CoV-2 challenge. Of note, RBD adjuvanted with AH:CpG elicited levels of neutralizing Abs comparable to the clinical-grade BNT162b2 Spike mRNA vaccine.

The translational relevance of our findings is also highlighted by the synergistic activation of human PBMCs from older individuals upon stimulation with AH:CpG. Overall, our results expand upon recent preclinical and clinical studies on the enhanced immunogenicity of Spike formulated with AH:CpG by showing that a vaccine composed of RBD and AH:CpG can also induce a robust anti-SARS-CoV-2 immune response across different age groups. Since an RBD antigen is amenable to high-yield manufacturability (*52-54*), our study also supports the development of RBD formulated with AH:CpG as an affordable and accessible vaccine.

Among various AH:PRR agonist formulations, AH:CpG elicited the highest immune responses in both young and aged mice. We observed that vaccine immune responses were generally lower in aged mice than in young adult mice, even in the group receiving RBD formulated with AH:CpG. While the lower levels of anti-RBD Abs observed in aged mice are likely sufficient for protection, we found that an additional booster dose in the aged overcame the observed age-dependent reductions in vaccine response and protected aged mice from SARS-CoV-2 challenge. We employed AH, which has been used for >90 years with a firmly established record of safety and efficacy (*32*) and AS01B, which recently demonstrated excellent adjuvant effects among elderly humans (*33, 34*), as “benchmarking” adjuvants to compare the exploratory adjuvanted formulations with more established adjuvants. In this context, we demonstrated that the AH:CpG adjuvanted vaccine was superior to a vaccine adjuvanted only with AH and was non-inferior to AS01B. In the context of the aged mice prime-boost setting, AH:CpG-adjuvanted SARS-CoV-2 RBD significantly outperformed AS01B with respect to functional anti-RBD inhibition (Geometric mean (GM) with SD, 57±2% *vs*. 14±3%) and neutralizing Abs titers (2344 ± 7 *vs.* 117± 4).

In this study, AH:CpG dramatically enhanced vaccine immune responses compared to vaccines adjuvanted with AH or CpG alone in both young and aged mice. AH:PRR agonist formulations have shown promising adjuvanticity in preclinical models, and AS04 (a formulation of aluminum salts and MPLA) is employed in several licensed vaccines (*32*). While the precise mechanism of action of AH:PRR agonist formulations has not been completely uncovered and is potentially influenced by the degree of adsorption of PRR ligands onto AH, the effects of these formulations are at least in part mediated by enhanced activation of innate immune cells at the injection site (*31, 55*). In our murine model, we also show that AH:CpG and CpG alone induce comparable proinflammatory gene expression profiles in dLNs. To gain additional mechanistic insight and increase the translational relevance of our findings, we tested the activity of AH:CpG on human PBMCs isolated from young adults and older individuals and found that this adjuvant formulation synergistically enhances cytokine and chemokine production compared to AH or CpG. These results might be explained by either 1) synergistic activation by AH and CpG of distinct molecular pathways, and/or 2) adsorption of CpG onto AH leading to the formation of macromolecular complexes that are more efficiently internalized and/or lead to enhanced TLR9 activation. Further work is required to define the underlying molecular mechanism of action of AH:CpG *in vivo* and *in vitro*.

The rationale for use of a synthetic TLR9 agonist CpG as an adjuvant for SARS-CoV-2 subunit vaccine is multi-fold. First, CpG has been used as a vaccine adjuvant in licensed vaccines with well-known mechanisms, substantial safety data, and confirmed effectiveness (*56, 57*). Second, CpG has demonstrated adjuvant effects in elderly populations. CpG enhanced vaccinal antigen immunogenicity in aged mouse and porcine models (*58-63*). Several human trials demonstrated that older individuals had a higher seroprotection rate when immunized with the CpG-adjuvanted hepatitis B vaccine compared to the conventional alum-adjuvanted vaccine (*64, 65*). Finally, AH:CpG-adjuvanted SARS-CoV-2 Spike vaccines have demonstrated safety, immunogenicity, and efficacy in several young adult animal models (*51, 66, 67*), and in a human clinical study involving an older population (*12*). Furthermore, Biological E has recently completed early phase (1 and 2) trials of a AH:CpG-adjuvanted SARS-CoV-2 RBD protein vaccine (trial # CTRI/2020/11/029032) which was intended for low- and middle-income countries, and are currently advancing through manufacturing and clinical development through a large-scale phase 3 trial in India (*17*). CpG is classified into 4 major classes, with distinct activation profiles of human cells (*68*). Class B CpG-1018 has been extensively evaluated in clinical trials. We observed that CpG-1018 and the class C CpG-2395 formulated with AH elicit comparable levels of neutralizing Abs, resulting in adjuvanted RBD formulations that were both non-inferior to the clinical-grade BNT162b2 Spike mRNA vaccine. Studies of TLR7/8 agonists as precision adjuvants with robust activity in early life (*69*), including in enhancing Spike immunogenicity in the young (*70*), further support the use of adjuvants to enhance vaccine immunogenicity in target populations. Together, and in light of our results in the older individuals, these studies suggest that precision adjuvant approaches hold substantial promise to generate scalable adjuvanted SARS-CoV-2 vaccine formulations that do not require freezing and afford robust protection to vulnerable populations across the lifespan.

Our study features several strengths, including (a) defining a combination adjuvantation system based on the common AH backbone that demonstrated mathematical synergy in its ability to activate human mononuclear cells; (b) accounting for age-specific immunity that can play major roles in vaccine immunogenicity and is often overlooked in vaccine discovery; (c) accounting for species-specificity by assessing the activity of the adjuvant formulation in human PBMCs *in vitro* and in mice *in vivo*; (d) testing the ability of the adjuvanted formulation to protect in a SARS-CoV-2 challenge model; and (e) benchmarking to the authorized BNT162b2 Spike mRNA vaccine to place our studies in context. As with any research our study also has some limitations, including that (a) we performed *in vivo* analysis only in mice, establishing the need for future translational research in additional animal models and humans and (b) all adjuvants/antigens were compared in single dose and further analysis should be performed in multiple doses to evaluate both efficacy and reactogenicity. Nevertheless, since we used standard doses of adjuvants/antigens in mouse systems (e.g., 1/30 and 1/18th of the human dose for CpG (*12*) and BNT162b2 (*3*) respectively, to compare the CpG-adjuvanted RBD subunit vaccine to the mRNA vaccine), it should be underscored that the results in this study hold promising value from a translational perspective.

Recently, several SARS-CoV-2 variants of concern have emerged harboring mutations in the RBD region and showing various degrees of reduced neutralization by serum samples obtained from convalescent or vaccinated individuals (*38-40*). It is likely that booster doses that account for mutations in the Spike protein will be required in order to achieve complete immunity against such variants (*71*). Several vaccines composed of multiple protein antigens adsorbed onto aluminum salts alone or co-formulated with MPLA have been produced (*55, 72*). We speculate that an AH:CpG-adjuvanted coronavirus vaccine formulation incorporating RBD proteins from different SARS-CoV-2 strains (and potentially other coronaviruses) may promote cross-strain protective immunity.

Overall, the current study aimed to evaluate an optimal adjuvant formulation to improve the protective response of RBD-based subunit vaccines in the elderly population, which is otherwise reduced as an effect of aging. We show that an AH:CpG adjuvant formulation induces potent anti-RBD responses in both young and aged mice and overcomes both the poor immunogenicity of the antigen and impaired immune responses in the aged. We discovered unique immunological properties of the AH:CpG adjuvant formulation that demonstrated synergistic enhancement of the production of multiple cytokines and chemokines from human adult and elderly PBMCs *in vitro*. These data indicate that formulating RBD with AH:CpG represents a promising approach to develop a practical (e.g., not requiring low temperature storage), scalable, effective, and affordable vaccine that may be effective across multiple age groups and could potentially incorporate multiple RBD proteins to achieve cross-strain protection.

## MATERIALS AND METHODS

### Study design

The aim of this study was to assess optimal combinations of RBD antigen and adjuvants in pre-clinical models that take age-dependent vaccine immune responses and COVID-19 susceptibility into account. To this end, we used age-specific mouse *in vivo* and human *in vitro* models. Sample size and age criteria was chosen empirically based on results of previous studies. Mouse experiments aimed to include in total 10 mice per group and were combined from two individual experiments. Mice were randomly assigned to different treatment groups. In order to assess the translational relevance and potential mechanism of an adjuvant formulation, we designed human *in vitro* study with peripheral blood collected from healthy young adults, aged 18–40 y (n = 6), and older participants, aged ≥ 65 years (n = 6), with approval from the Ethics Committee of the Boston Children’s Hospital (protocol number X07-05-0223) and Institutional Review Board of Brigham and Women’s Hospital, Boston (protocol number 2013P002473). All participants signed an informed consent form prior to enrollment. Investigators were not blinded. No data outliers were excluded.

### Animals

Female, 3 months old BALB/c mice were purchased from Jackson Laboratory (Bar Harbor, ME). Female, 12-13 months old BALB/c mice purchased from Taconic Biosciences (Germantown, NY) were used for aged mice experiments. Mice were housed under specific pathogen-free conditions at Boston Children’s Hospital, and all the procedures were approved under the Institutional Animal Care and Use Committee (IACUC) and operated under the supervision of the Department of Animal Resources at Children’s Hospital (ARCH) (Protocol number 19-02-3897R). At the University of Maryland School of Medicine, mice were housed in a biosafety level 3 (BSL3) facility for all SARS-CoV-2 infections with all the procedures approved under the IACUC (Protocol number #1120004) to MBF.

### SARS-CoV-2 Spike and RBD expression and purification

Full length SARS-CoV-2 Spike glycoprotein (M1-Q1208, GenBank MN90894) and RBD constructs (amino acid residues R319-K529, GenBank MN975262.1), both with an HRV3C protease cleavage site, a TwinStrepTag and an 8XHisTag at C-terminus were obtained from Barney S. Graham (NIH Vaccine Research Center) and Aaron G. Schmidt (Ragon Institute), respectively. These mammalian expression vectors were used to transfect Expi293F suspension cells (Thermo Fisher) using polyethylenimine (Polysciences). Cells were allowed to grow in 37°C, 8% CO_2_ for additional 5 days before harvesting for purification. Protein was purified in a PBS buffer (pH 7.4) from filtered supernatants by using either StrepTactin resin (IBA) or Cobalt-TALON resin (Takara). Affinity tags were cleaved off from eluted protein samples by HRV 3C protease, and tag removed proteins were further purified by size-exclusion chromatography using a Superose 6 10/300 column (Cytiva) for full length Spike and a Superdex 75 10/300 Increase column (Cytiva) for RBD domain in a PBS buffer (pH 7.4).

### Adjuvants and immunization

The adjuvants and their doses used were: Alhydrogel adjuvant 2% (100 µg), 2’3’-cGAMP (10 µg), Poly (I:C) HMW (50 µg), CpG-ODN 2395 (50 µg), AddaS03 (25 µL) (all from InvivoGen, San Diego, CA), CpG-ODN 1018 (50 µg, 5’ TGA CTG TGA ACG TTC GAG ATG A 3’) (Integrated DNA Technologies, Coralville, IA), PHAD (50 µg) (Avanti Polar Lipids, Alabaster, AL), and AS01B (40 µL) (obtained from the Shingrix vaccine, GSK Biologicals SA, Belgium). Mice were injected with 10 µg of recombinant monomeric SARS-CoV-2 RBD protein, with or without adjuvants. Each PRR agonist was formulated with and without aluminum hydroxide. Mock treatment mice received phosphate-buffered saline (PBS) alone. BNT162b2 Spike mRNA vaccine (Pfizer-BioNTech) was obtained as residual volumes in used vials from the Boston Children’s Hospital employee vaccine clinic, strictly using material that would only otherwise be discarded, and was used within 6 hours from the time of reconstitution. BNT162b2 suspension (100 µg/mL) was diluted 1:3 in PBS, and 50 µL (1.67 µg) was injected. Injections (50 µL) were administered intramuscularly in the caudal thigh on Days -0, -14 (both age groups), and Day 28 (aged mice only, where relevant). Blood samples were collected 2 weeks post-immunization.

### ELISA

RBD- and Spike-specific antibody levels were quantified in serum samples by ELISA by modification of a previously described protocol(*73*). Briefly, high-binding flat-bottom 96-well plates (Corning, NY) were coated with 50 ng/well RBD or 25 ng/well Spike and incubated overnight at 4 °C. Plates were washed with 0.05% Tween 20 PBS and blocked with 1% BSA PBS for 1 h at room temperature (RT). Serum samples were serially diluted 4-fold from 1:100 up to 1:1.05E8 and then incubated for 2 hours at RT. Plates were washed three times and incubated for 1 hour at RT with HRP-conjugated anti-mouse IgG, IgG1, IgG2a, or IgG2c (Southern Biotech). Plates were washed five times and developed with tetramethylbenzidine (1-Step Ultra TMB-ELISA Substrate Solution, ThermoFisher, for RBD-ELISA, and BD OptEIA Substrate Solution, BD Biosciences, for Spike ELISA) for 5 min, then stopped with 2 N H_2_SO_4_. Optical densities (ODs) were read at 450 nm with SpectraMax iD3 microplate reader (Molecular Devices). End-point titers were calculated as the dilution that emitted an optical density exceeding a 3× background. An arbitrary value of 50 was assigned to the samples with OD values below the limit of detection for which it was not possible to interpolate the titer.

### hACE2/RBD inhibition assay

The hACE2/RBD inhibition assay employed a modification of a previously published protocol(*74*). Briefly, high-binding flat-bottom 96-well plates (Corning, NY) were coated with 100 ng/well recombinant human ACE2 (hACE2) (Sigma-Aldrich) in PBS, incubated overnight at 4°C, washed three times with 0.05% Tween 20 PBS, and blocked with 1% BSA PBS for 1 hour at RT. Each serum sample was diluted 1:160, pre-incubated with 3 ng of RBD-Fc in 1% BSA PBS for 1 hour at RT, and then transferred to the hACE2-coated plate. RBD-Fc without pre-incubation with serum samples was added as a positive control, and 1% BSA PBS without serum pre-incubation was added as a negative control. Plates were then washed three times and incubated with HRP-conjugated anti-human IgG Fc (Southern Biotech) for 1 hour at RT. Plates were washed five times and developed with tetramethylbenzidine (BD OptEIA Substrate Solution, BD Biosciences) for 5 min, then stopped with 2 N H_2_SO_4_. The optical density was read at 450 nm with SpectraMax iD3 microplate reader (Molecular Devices). Percentage inhibition of RBD binding to hACE2 was calculated with the following formula: Inhibition (%) = [1 – (Sample OD value – Negative Control OD value)/(Positive Control OD value – Negative Control OD value)] x 100.

### SARS-CoV-2 neutralization titer determination

All serum samples were heat-inactivated at 56°C for 30 min to remove complement and allowed to equilibrate to RT prior to processing for neutralization titer. Samples were diluted in duplicate to an initial dilution of 1:5 or 1:10 followed by 1:2 serial dilutions (vaccinated sample), resulting in a 12-dilution series with each well containing 100 µL. All dilutions were performed in DMEM (Quality Biological), supplemented with 10% (v/v) fetal bovine serum (heat-inactivated, Sigma), 1% (v/v) penicillin/streptomycin (Gemini Bio-products) and 1% (v/v) L-glutamine (2 mM final concentration, Gibco). Dilution plates were then transported into the BSL-3 laboratory and 100µL of diluted SARS-CoV-2 (WA-1, courtesy of Dr. Natalie Thornburg/CDC) inoculum was added to each well to result in a multiplicity of infection (MOI) of 0.01 upon transfer to titering plates. A non-treated, virus-only control and mock infection control were included on every plate. The sample/virus mixture was then incubated at 37°C (5.0% CO_2_) for 1 hour before transferring to 96-well titer plates with confluent VeroE6 cells. Titer plates were incubated at 37°C (5.0% CO_2_) for 72 hours, followed by CPE determination for each well in the plate. The first sample dilution to show CPE was reported as the minimum sample dilution required to neutralize >99% of the concentration of SARS-CoV-2 tested (NT99).

### Pseudovirus neutralization assay

The SARS-CoV-2 pseudoviruses expressing a luciferase reporter gene were generated in an approach similar to as described previously (*75, 76*). Briefly, the packaging plasmid psPAX2 (AIDS Resource and Reagent Program), luciferase reporter plasmid pLenti-CMV Puro-Luc (Addgene), and spike protein expressing pcDNA3.1-SARSCoV-2 SΔCT of variants were co-transfected into HEK293T cells by lipofectamine 2000 (ThermoFisher). Pseudoviruses of SARS-CoV-2 variants were generated by using WA1/2020 strain (Wuhan/WIV04/2019, GISAID accession ID: EPI_ISL_402124), B.1.1.7 variant (GISAID accession ID: EPI_ISL_601443), or B.1.351 variant (GISAID accession ID: EPI_ISL_712096). The supernatants containing the pseudotype viruses were collected 48 h post-transfection, which were purified by centrifugation and filtration with 0.45 µm filter. To determine the neutralization activity of the plasma or serum samples from participants, HEK293T-hACE2 cells were seeded in 96-well tissue culture plates at a density of 1.75 x 10^4^ cells/well overnight. Three-fold serial dilutions of heat inactivated serum or plasma samples were prepared and mixed with 50 µL of pseudovirus. The mixture was incubated at 37°C for 1 h before adding to HEK293T-hACE2 cells. 48 h after infection, cells were lysed in Steady-Glo Luciferase Assay (Promega) according to the manufacturer’s instructions. SARS-CoV-2 neutralization titers were defined as the sample dilution at which a 50% reduction in relative light unit (RLU) was observed relative to the average of the virus control wells.

### Splenocyte restimulation assay

Immunized mice were sacrificed 2 weeks after the final immunization, and spleens were collected. To isolate splenocytes, spleens were mashed through a 70 µm cell strainer, and the resulting cell suspensions were washed with PBS and incubated with 2 mL of ACK lysis buffer (Gibco) for 2 minutes at RT to lyse erythrocytes. Splenocytes were washed again with PBS and plated in flat-bottom 96-well plates (2 x 10^6^ cells per well). Then, SARS-CoV-2 Spike peptides (PepTivator SARS-CoV-2 Prot_S, Miltenyi Biotec) were added at a final concentration of 0.6 nmol/ml in the presence of 1 μ (total cell culture volume, 200 µL per well). After 24 (for IL-2 and IL-4) and 96 (for IFNγ) hours, supernatants were harvested, and cytokine levels were measured by ELISA (Invitrogen) according to the manufacturer’s protocol.

### SARS-CoV-2 mouse challenge study

Mice were anesthetized by intraperitoneal injection 50 μL of a mix of xylazine (0.38 mg/mouse) and ketamine (1.3 mg/mouse) diluted in PBS. Mice were then intranasally inoculated with 1 x 10^3^ PFU of mouse-adapted SARS-CoV-2 (MA10, courtesy of Dr. Ralph Baric (UNC)) in 50 μ divided between nares(*35*). Different doses of SARS-CoV-2 were used where indicated. Challenged mice were weighed on the day of infection and daily for up to 4 days post-infection. At 4-day post-infection, mice were sacrificed, and lungs were harvested to determine virus titer by a plaque assay and prepared for histological scoring.

### SARS-CoV-2 plaque assay

SARS-CoV-2 lung titers were quantified by homogenizing harvested lungs in PBS (Quality Biological Inc.) using 1.0 mm glass beads (Sigma Aldrich) and a Beadruptor (Omni International Inc.). Homogenates were added to Vero E6 cells and SARS-CoV-2 virus titers determined by counting plaque-forming units (pfu) using a 6-point dilution curve.

### Histopathology analysis

Slides were prepared as 5-μm sections and stained with hematoxylin and eosin. A pathologist was blinded to information identifying the treatment groups and fields were examined by light microscopy and analyzed. The severity of interstitial inflammation was evaluated and converted to a score of 0-4 with 0 being no inflammation and 4 being most severe. Interstitial inflammation was evaluated for the number of neutrophils present in the interstitial space as well as the extent of neutrophilic apoptosis. Once scoring was complete, scores for each group were averaged and the standard deviation for the scoring was computed.

### Mouse *in vivo* LNs gene expression analysis by quantitative real-time PCR array

Mice were subcutaneously injected on Day 0 with the indicated treatments in a volume of 50 µL on each side of the back (one side for the compound and the contralateral side for saline of vehicle control). Twenty-four hours post-injection, draining (brachial) LNs were collected for subsequent analysis. LNs were transferred to a beadbeater and homogenized in TRI Reagent (Zymo Research). Samples were then centrifuged, and the clear supernatant was transferred to a new tube for subsequent RNA isolation. RNA was isolated from TRI Reagent samples using phenol-chloroform extraction or column-based extraction systems (Direct-zol RNA Miniprep, Zymo Research) according to the manufacturer’s protocol. RNA concentration and purity (260/280 and 260/230 ratios) were measured by NanoDrop (ThermoFisher Scientific). cDNA was prepared from RNA with RT^2^ First Strand Kit, according to the manufacturer’s instructions (Qiagen). cDNA was quantified using 96-well PCR array analysis on a PAMM-150ZA plate (Cytokines & Chemokines) and PAMM-016ZA plate (Type I Interferon Response) (both Qiagen). Quantitative real time-PCR (QRT-PCR) was run on a 7300 real-time PCR system (Applied Biosystems – Life Technologies, Carlsbad, CA). mRNA levels were normalized to 3 housekeeping genes: *Actb*, *Gapdh*, and *Gusb*. Relative quantification of gene expression was ΔΔ*Ct* (relative expression over PBS treatment group).

### Human PBMC isolation

PBMCs were isolated based previously described protocols (*77*). Briefly, heparinized whole blood was centrifuged at 500 g for 10 min, then the upper layer of platelet-rich plasma was removed. Plasma was centrifuged at 3000 g for 10 min, and platelet-poor plasma (PPP) was collected and stored on ice. The remaining blood was reconstituted to its original volume with heparinized DPBS and layered on Ficoll-Paque gradients (Cytiva) in Accuspin tubes (Sigma-Aldrich). PBMCs were collected after centrifugation and washed twice with PBS.

### Human PBMCs stimulation

PBMCs were resuspended at a concentration of 200,000 cells per well in a 96-well U-bottom plate (Corning) in 200 µL RPMI 1640 media (Gibco) supplemented with 10% autologous PPP, 100 IU/mL penicillin, 100 µg/mL streptomycin, and 2 mM L-glutamine. PBMCs were incubated for 24 hrs at 37°C in a humidified incubator at 5% CO_2_ with indicated treatments. After culture, plates were centrifuged at 500 g and supernatants were removed by pipetting without disturbing the cell pellet. Cytokine expression profiles in cell culture supernatants were measured using customized Milliplex human cytokine magnetic bead panels (Milliplex). Assays were analyzed on the Luminex FLEXMAP 3D employing xPONENT software (Luminex) and Millipore Milliplex Analyst. Cytokine measurements were excluded from analysis if fewer than 30 beads were recovered. Synergy was evaluated using the Loewe definition of additivity, with D > 1 indicating antagonism, D = 1 additivity, and D < 1 synergy (*78*). In order to fit regression curves more closely to the data, higher concentrations were excluded from linear regressions when calculating D values if the cytokine levels plateaued or decreased.

### Statistical analysis

Statistical analyses were performed using Prism v9.0.2 (GraphPad Software) and R software environment v4.0.4. *P* values < 0.05 were considered significant. Data were analyzed by one- or two-way ANOVAs followed by post-hoc Tukey’s test or Dunnett’s test for multiple comparisons. Non-normally distributed data were log-transformed. In the animal experiences, time to event were analyzed using Kaplan-Meier estimates and compared across groups using the Log-rank test. For human in vitro PBMC assay, unpaired Mann-Whitney tests were applied at each concentration. We conducted gene expression analyses with R 4.0.4 using packages ‘ggplot2’, ‘dplyr’, and ‘MASS’ for the transcript abundance determination of gene arrays in each group. We log-transformed data before performing principal component analysis (PCA) and unsupervised hierarchical clustering using R packages ‘prcomp’ and ‘pheatmap’ respectively. We analyzed the differential gene expression using generalized linear models (GLMs) with treatment and age as fixed effects. We then enriched the differentially expressed genes using the blood transcriptional module method based on an existing protocol (Li et al., 2013-PMC: 24336226).

## ACKNOWLEDGEMENTS

We thank the members of the BCH *Precision Vaccine Program* for helpful discussions. We thank Drs. Kevin Churchwell, Gary Fleisher, David Williams, and Mr. August Cervini for their support of the *Precision Vaccines Program*. We thank Dr. Barney S. Graham (NIH Vaccine Research Center) for generously providing the plasmid for pre-fusion stabilized SARS-CoV-2 Spike trimer. We thank Dr. Ralph Baric for providing the SARS-CoV-2/MA10 virus. We thank the pharmacists of Boston Children’s Hospital for their efforts to maximize the use of SARS-CoV-2 vaccines by saving leftover or overfill of otherwise to be discarded vaccine vials. D.J.D. would like to thank Ms. Siobhan McHugh, Ms. Geneva Boyer, Mrs. Lucy Conetta and the staff of Lucy’s Daycare, the staff the YMCA of Greater Boston, Bridging Independent Living Together (BILT), Inc., and the Boston Public Schools for childcare and educational support during the COVID-19 pandemic.

## AUTHOR CONTRIBUTIONS

EN and FB conceived, designed, performed, analyzed the experiments and wrote the paper; TRO, YS, BB, MDL, KC, MMe, SDG, and SVH performed *in vitro* and/or *in vivo* experiments and their analysis; JC and JD-A performed the analysis of the qPCR data; KS, AZX, HSS, SDP, TMC, JF, BMH, AGS expressed and purified SARS-CoV-2 RBD and Spike; MEM, REH, CD, SMW, RMJ, HLH, RM, AB and MBF performed and analyzed SARS-CoV-2 neutralization experiments and mouse challenge study; AC, JY and DHB performed and analyzed pseudovirus neutralization experiments; ACS and LRB contributed to the human elderly *in vitro* analysis; MEB, PJH, and US edited and critically reviewed the manuscript; RKE and IZ contributed to the design of experiments; AO provided design feedback and contributed to the statistical analysis; OL and DJD concexived the project, designed the experiments, supervised the study and wrote the paper.

## DATA AND MATERIALS AVAILABILITY

All data are available in the main text or the supplementary materials

## SUPPLEMENTARY MATERIALS

**Supplementary Figure 1.**
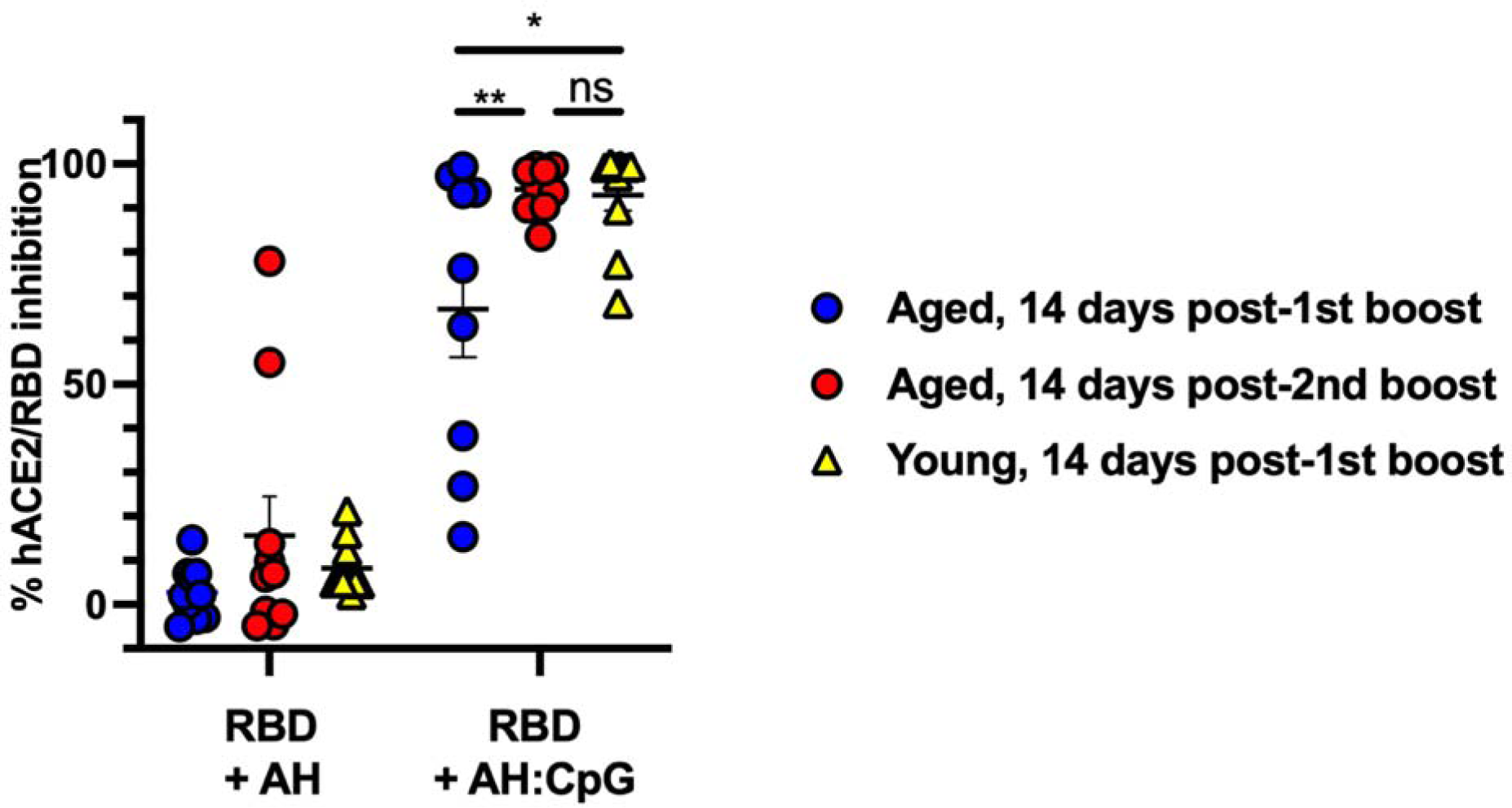
Booster dose of AH:CpG formulation enhances hACE2/RBD inhibition in aged mice. Young adult, 3-month-old BALB/c mice were immunized IM on Days 0 and 14, and aged, 14-month-old BALB/c mice were immunized IM on Days 0, 14, and 28 with 10 µg of monomeric SARS-CoV-2 RBD protein with the indicated adjuvants. Serum samples were collected and analyzed on Day 28 prior to the 2nd boost, and Day 42. hACE2/RBD inhibition rate was assessed. N = 9-10 animals per group. Data were combined from two individual experiments and analyzed by one-way ANOVA followed by post-hoc Tukey’s test for multiple comparisons. Each dot represents individual results. Horizontal bars demonstrate mean plus SEM. ns: not significant, \**P* <0.05, \*\**P* <0.01. AH, aluminum hydroxide.

**Supplementary Figure 2.**
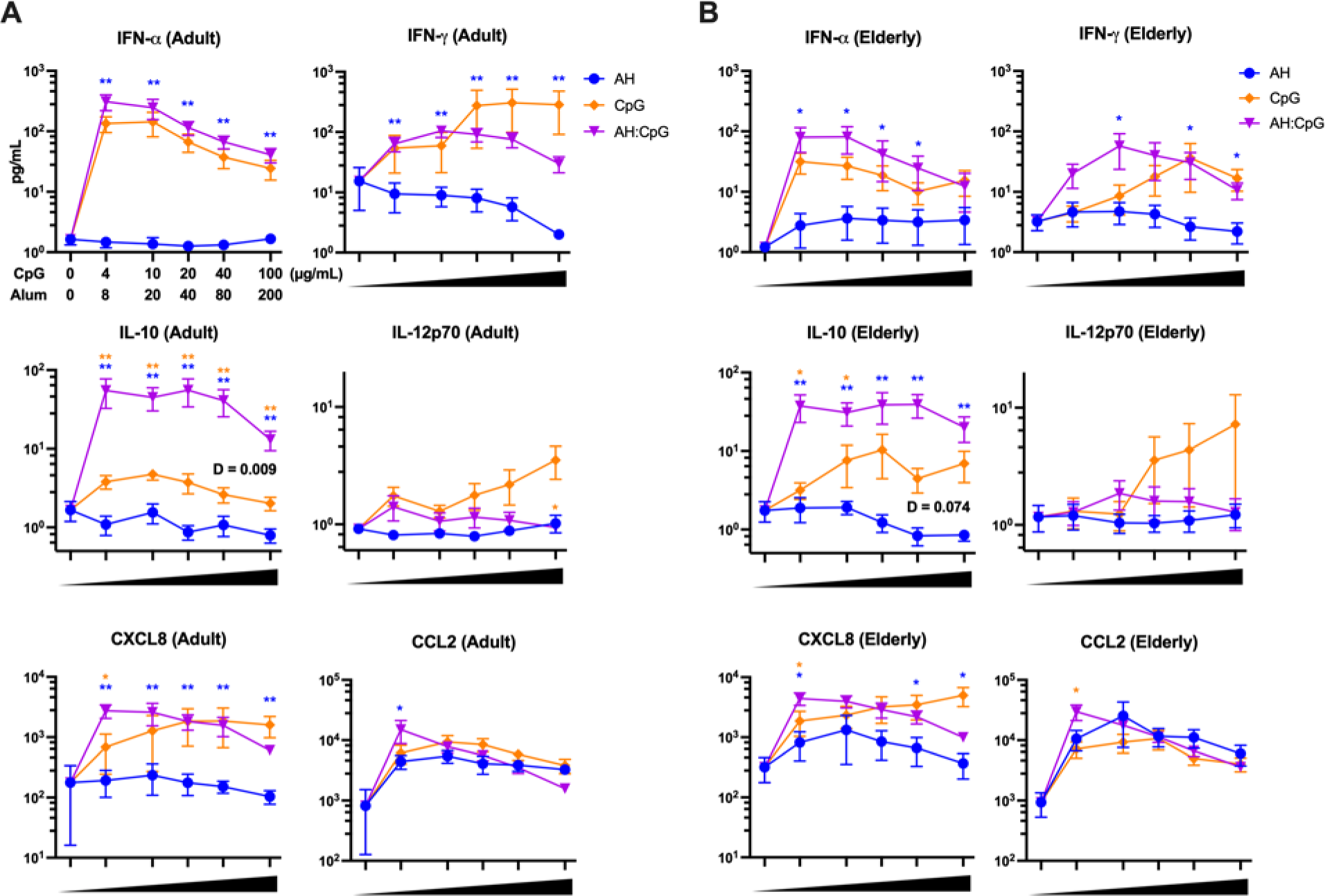
AH and CpG synergistically induce cytokine and chemokine production by human young adult and elderly PBMCs. Human PBMCs collected from young adults (**A**) and elderly individuals (**B**) were cultured *in vitro* for 24 hrs with CpG alone (4, 10, 20, 40, and 100 μg/mL), aluminum hydroxide (AH) alone (8, 20, 40, 80, and 200 μg/mL), or combinations of each. Supernatants were collected for multiplexing bead array. N=6 per age group. Unpaired Mann-Whitney tests were applied at each concentration. Level of synergy was calculated using an adapted Loewe definition of additivity (D <1: synergy, D=1: additivity, D >1: antagonism). D value was not calculated if the concentration-dependent cytokine level did not fit a linear regression curve. Blue and yellow colored asterisks indicate comparisons of AH:CpG formulation to AH and CpG alone groups, respectively. Results represent mean ± SEM. \**P* <0.05, \*\**P* <0.01.

